# The effect of preproenkephalin gene deletion on heart function in mice

**DOI:** 10.64898/2026.10.06.756947

**Authors:** Klaudia Misiołek, Łukasz Szumiec, Krzysztof Jasiński, Katarzyna Kalita, Maria Kaczmarczyk-Jarosz, Magdalena Chrószcz, Władysław P. Węglarz, Barbara Ziółkowska, Jan Rodriguez Parkitna

## Abstract

Enkephalins, the endogenous ligands of the µ- and δ-opioid receptors, are abundantly expressed in the nervous system and peripheral organs, including the heart. Accumulating evidence suggests that they participate in cardiac development and act as autocrine and paracrine modulators of heart activity. Here, we investigate the role of enkephalins in cardiac physiology using a newly generated mouse strain with a complete ablation of the preproenkephalin (*Penk*) gene. We find that, although adult mutant mice weighed less, the mutation caused no apparent changes in behavioral phenotype, no increase in anxiety-like behaviors, and only a mild reduction in locomotor activity. There was no effect of *Penk* deletion on heart weight or size. A comprehensive analysis of left ventricular function using MRI revealed normal basal cardiac performance, except for a significant reduction in the peak fill and ejection rates. There were no significant effects of *Penk* deletion on left ventricular function after β_1_-adrenergic stimulation with dobutamine. At the same time, we observed the expected significant differences in heart size and function between male and female mice. In conclusion, our data show that enkephalins are not essential for normal heart development and have a moderate impact on adult heart function under basal conditions.

## 1 Introduction

Enkephalins are ubiquitous opioid peptides abundant in the central and enteric nervous systems, the adrenal medulla, gonads, and the cardiac tissue [1–4]. They are derived from the protein preproenkephalin (Penk), the product of the *Penk* gene, which is cleaved into peptides of different lengths that share the common N-terminal sequence Tyr-Gly-Gly-Phe. Enkephalins include the pentapeptides Met- and Leu-enkephalin, and several longer peptides containing the Met-enkephalin sequence [2]. They are endogenous ligands of the µ-, δ-, and, to a lesser extent, κ-opioid receptors, with different Penk-derived peptides exhibiting distinct receptor affinities. Met- and Leu-enkephalin have the highest affinity for δ-opioid receptors and slightly lower affinity for µ-opioid receptors [2]. Nevertheless, in the latter case, the reported K_d_ values are the same or lower than those for β-endorphin [2]. Activation of opioid receptors on neuronal cells by enkephalins inhibits their activity by causing hyperpolarization and blocking calcium influx [5]. Enkephalins have been extensively studied for their roles in nociception, the stress response, the activity of the reward system, and a wide range of other brain functions [2,4]. Conversely, the functions of *Penk*-derived peptides outside the nervous system have received less attention.

Cardiac tissue contains large amounts of *Penk* mRNA, comparable to or even higher than those in the brain [1,3,6]. Cardiomyocytes synthesize and process enkephalins [3,7], and the release of Met-enkephalin, as well as longer Penk-derived peptides, from cardiac cells has been described in cell cultures and in isolated rat hearts [7,8]. Enkephalins have been proposed to exert paracrine and autocrine effects within the heart, acting both directly on cardiomyocytes and on autonomic nerve endings [9,10]. This notion is supported by the presence of the preferential enkephalin target, the δ-opioid receptor, and its mRNA in the cardiac tissue [1,11]. Enkephalins acting via the δ-opioid receptors were reported to reduce cardiac cell contractility and attenuate the positive inotropic effect of β-adrenergic receptor stimulation in isolated cardiomyocytes and rat hearts [10,12,13]. They were also shown to regulate heart rate by modulating sympathetic and parasympathetic inputs to the sinoatrial node, most likely acting on prejunctional opioid receptors [14–16]. Moreover, Met-enkephalin plays a protective role in cardiac ischemia and mediates ischemic preconditioning. Enkephalins are released by the heart during an ischemic episode and limit infarct development in a δ-opioid receptor-dependent fashion (reviewed in[9]). Apart from ischemia, cardiac enkephalins or *Penk* mRNA are also up-regulated in ageing, hypertension, and cardiomyopathy [9,17]. One of the Penk-derived peptides released into the bloodstream, Proenkephalin-A, has been proposed to be a marker of heart failure severity in humans, although its clinical relevance remains unclear [18,19].

Enkephalins have also been implicated in heart development, where Met-enkephalin was reported to be an essential inhibitory growth factor [20]. It was demonstrated that rats treated with the opioid antagonist naltrexone either prenatally or on the first postnatal day developed larger hearts than controls [21,22]. The effect on cardiac growth was shown to be specific to Met-enkephalin, and was not reproduced by other opioid peptides or exogenous agonists of the classical (µ, δ, and κ) opioid receptors [20,23]. The inhibition of cardiac growth was linked to activation of a non-canonical opioid receptor, zeta (ζ), located in the nuclear envelope, which, upon binding Met-enkephalin, inhibits DNA synthesis and cell proliferation [20,22]. Here, we investigate the contribution of Penk-derived peptides to cardiac development and function using a novel mouse strain with complete *Penk* gene inactivation.

## 2 Materials and Methods

### 2.1 Penk KO mice

The *Penk^tm1Nfmol^* (*Penk* KO) strain was generated in two steps. First, a floxed variant of the *Penk* gene was generated as a service by Cyagen. LoxP sites were introduced flanking the second exon, in the 5’ and 3’ UTR sequences (Fig. 1A). The targeting vector was used for homologous recombination in the 5C11 mouse embryonic stem cell line and the generation of founders. Initial characterization of the strain revealed that the insertion of loxP sites had significantly reduced *Penk* gene expression, prompting a second step to convert the line from a conditional floxed allele into a full germline knockout. Mice with the floxed *Penk* gene were crossed with animals from the B6.Cg-Tg(Syn1-cre)671Jxm/J strain [24], which led to complete deletion in the *Penk* allele in the offspring. The entire protein-coding region was removed in the resulting *Penk^tm1Nfmol^* strain. This strain was bred at the animal facility of the Maj Institute of Pharmacology of the Polish Academy of Sciences as a congenic line with C57BL/6J for 6 generations before experimental procedures began. Genotype distribution among *Penk^tm1Nfmol^* offspring followed expected Mendelian ratios.

**Figure 1.**
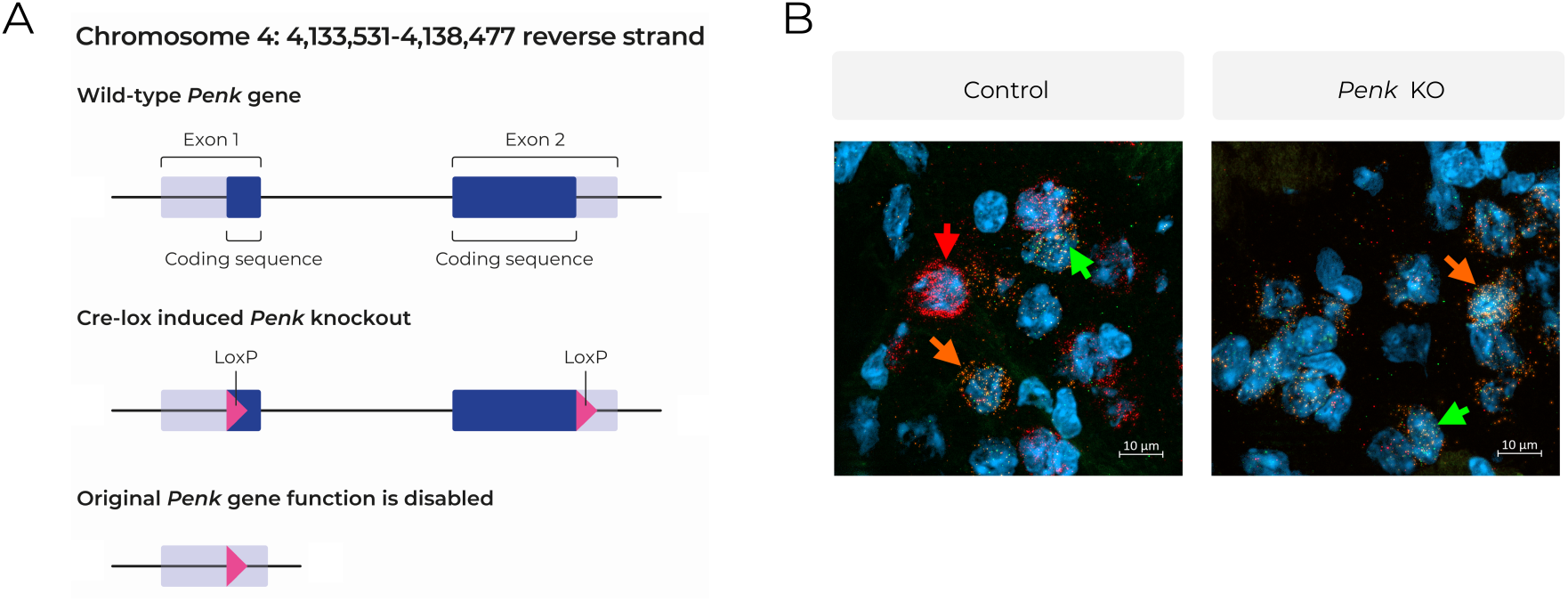
Generation and validation of proenkephalin knockout (*Penk* KO) mice. (A) Schematic representation of the genetic modification resulting in deletion of the *Penk* protein-encoding sequence. (B) Representative RNAscope images from mouse brain (dorsal striatum) of control and *Penk* KO mice. DAPI (blue), *Pdyn* (Atto550, orange), *Penk* (Atto647, red), and *Oprd1* (Alexa Fluor 488, green) mRNA expression is shown. Arrows indicate example transcript sites; arrow color corresponds to the fluorophore described above. Scale bars are shown in each image. Control and *Penk* KO animals are shown in gray and blue, respectively.

Genotyping of *Penk* KO mice was performed by PCR using primers CTACAGGCGCGTTCTTCTCT and GGATGGAGCAGGTAGTGG, and the following conditions: initial denaturation 3’ at 95°C, followed by 40 cycles of 30 s denaturation at 95°C, primer annealing 30s at 58°C, and elongation 60s at 72°C. The reaction amplifies a 110 bp DNA product in the case of a KO allele and a 134 bp product in the case of a wild-type allele. Animals were genotyped at weaning and, for experimental mice, also post-mortem.

### 2.2 General Procedures

Sex of the animals was determined at weaning based on anogenital distance and the presence or absence of nipples. All behavioral tests were conducted during the light phase of the circadian cycle. Mice were group-housed (2–6 same-sex littermates per cage) in standard Plexiglas cages (325 × 170 mm) containing nesting material and wooden blocks, under controlled laboratory conditions (22 ± 2°C; 40–60% relative humidity; 12:12 hours light/dark cycle, lights on at 07:00). Food (Altromin Spezialfutter, cat. no. 1324, Germany) and water were available *ad libitum*. Behavioral testing was conducted under dim white light (5–10 lux) with infrared illumination. Mice were habituated to handling for 2–3 minutes daily over 3–5 consecutive days prior to testing. On each testing day, animals were acclimated to the experimenter and transferred to the testing room 30 minutes before the session to reduce novelty-induced stress. Behavioral activity was recorded using a Basler acA1300–60 gm camera (Basler, Germany) and analyzed using EthoVision XT v11.5 (Noldus Information Technology, The Netherlands) or BORIS v8.25 (Behavioral Observation Research Interactive Software; University of Turin, Italy). All procedures were approved by the 2nd Local Bioethics Committee in Kraków (protocol no. 27/2021) and were conducted in accordance with European Directive 2010/63/EU and Polish national regulations (Dz.U. 2015 poz. 266). Experimental design and reporting followed the ARRIVE guidelines. Details of experimental groups and animal numbers are provided in Supplementary Tables S1 and S2.

### 2.3 Exploratory behavior

Exploratory behavior was assessed using the open-field test. Mice were individually placed in the center of a white, rectangle arena (55 × 37.5 × 20.5 cm) with 0.5 cm of aspen bedding (ABEDD, Latvia), uniformly illuminated with dim white light (6–10 lux). Each animal was allowed to explore the arena freely for 30 minutes. A virtual central zone was defined as a square extending 5 cm inward from all walls to assess center-avoidance behavior. Behavioral activity was recorded and analyzed semi-automatically using EthoVision XT. The following parameters were quantified: total distance traveled [m], and percent of time spent in the central zone.

### 2.4 Social novelty preference

Social behavior was assessed using the social interaction test in the open field. Immediately following the assessment of exploratory behavior, a novel, age- and sex-matched conspecific was introduced into the same arena. Mice were allowed to interact freely for 10 minutes. Sessions were video-recorded and manually scored using BORIS software. Social interaction was defined as the cumulative time [s] the test mouse engaged in direct investigation or affiliative behaviors toward the stimulus animal, including sniffing (head, anogenital, or body), grooming, chasing, and mounting.

### 2.5 Anxiety-like behavior

We evaluated anxiety-like behavior using the light–dark box test. The apparatus consisted of two compartments (each 18 × 16 × 20 cm): a brightly lit box (400 lux) and a dark box (50 lux), connected via a small opening that allowed unrestricted movement between the two. Mice were placed in the dark compartment and allowed to freely explore the apparatus for 5 minutes. Behavior was recorded and analyzed semi-automatically using EthoVision XT. The following parameters were measured: latency to first entry into the light box [s], total time spent in the light box [s], total distance traveled [m] within the light box.

### 2.6 Anhedonia-like behavior

To assess anhedonia, we used a saccharin preference test based on a free-choice two-bottle paradigm. Twenty-four hours prior to testing, mice were individually housed in transparent cages (26 × 20 × 13.5 cm) containing standard bedding (ABEDD, Latvia), nesting material, a wooden chew block, and two 25-ml bottles filled with tap water to habituate the animals to the experimental setup. On the test day, both bottles were refilled, one with tap water and the other with a 0.1% (w/v) saccharin solution. Bottle position (left/right) was randomized across animals to control for potential side preference. Food was available *ad libitum* throughout the test. After 24 hours, fluid intake from each bottle was measured. The primary outcome measure was saccharin preference, calculated as: Saccharin Preference = (Saccharin intake [ml])/Total solution intake [ml]) × 100 %.

### 2.7 Heart weight

Mice were weighed using an electronic scale with an accuracy of 0.1 g. Following euthanasia by cervical dislocation, the heart was rapidly excised, placed on a paper towel for approximately 1 minute to allow excess blood to drain, and then weighed using an electronic scale with an accuracy of 0.1 mg. To normalize heart weight to body size, the left tibia was isolated and its length measured using a digital caliper with an accuracy of 0.01 mm. Two indices were calculated: heart weight to tibia length [mg/mm] and heart weight to body weight [mg/g].

### 2.8 RNA isolation and Penk qPCR

Immediately after weighing, the hearts were placed in 500 μl of RNAlater Stabilization Solution (Invitrogen, USA) for 1 hour at room temperature and then stored at 4°C. RNA was extracted and purified using the RNeasy Mini Kit (#74904, Qiagen) according to the manufacturer’s protocol. Reverse transcription was performed using the Omniscript RT Kit (Qiagen Inc., USA) following the manufacturer’s instructions. qPCR was performed using TaqMan probes targeting *Penk* (Mm01212875_m1) and *Hprt1* (Mm03024075_m1). Relative *Penk* transcript abundance was normalized to *Hprt* levels using 2^-ΔCycle threshold (Ct) calculation: Relative expression = 2^-(Ct*_Penk_ –* Ct*_Hprt_*).

### 2.9 Cardiovascular Magnetic Resonance Imaging (MRI)

Cardiac performance was assessed *in vivo* using a preclinical 9.4 T MRI scanner (BioSpec 94/20, Bruker BioSpin, Ettlingen, Germany) equipped with the BFG 102/60-S13 high-performance gradient insert and controlled via ParaVision 5.1 software. Radiofrequency (RF) transmission and signal reception were achieved using a quadrature birdcage resonator with a 36-mm bore diameter.

During the experiment, animals were anesthetized with isoflurane (1.7–2.0 vol%), in a carrier gas composed of oxygen and medical air at a 1:2 ratio. To ensure hemodynamic stability and prevent dehydration, a subcutaneous bolus of 0.9% NaCl (200 µl) was administered prior to positioning the animal in a supine orientation within a dedicated MRI-compatible cradle.

Physiological parameters were continuously monitored using an integrated monitoring system (SA Instruments, Stony Brook, NY, USA). Rectal temperature was continuously recorded and thermoregulation was maintained within the 36–37 °C range by means of a closed-loop water-heated pad. Respiratory rate was kept between 70 and 100 breaths/min by adjusting the isoflurane concentration, and heart rate (HR) was monitored continuously using a three-electrode ECG configuration.

Left ventricular (LV) function was assessed using seven contiguous short-axis slices, covering the entire ventricle from base to apex. Cine images were acquired as self-gated gradient-echo FLASH acquisition with retrospective respiratory and cardiac gating (IntraGate module, Bruker BioSpin) with the following parameters: field of view (FOV) = 30 × 30 mm^2^, matrix = 192 × 192, slice thickness = 1.0 mm, echo time (TE) = 1.3 ms, repetition time (TR) = 5.7 ms, flip angle = 19°, and 200 repetitions. Raw data were reconstructed into 20 cine frames per cardiac cycle using the standard IntraGate reconstruction algorithm.

Cardiac contractile reserve was evaluated using dobutamine stress MRI focused on a single mid-ventricular short-axis slice. To ensure direct comparability, identical acquisition parameters were used for both baseline and stress conditions: FOV = 30 × 30 mm^2^, matrix = 192 × 192, slice thickness = 1.0 mm, TE/TR = 1.5/4.5 ms, flip angle = 19°, and 400 repetitions. Dobutamine (Polfa Tarchomin, Warsaw, Poland) was administered intraperitoneally at a dose of 2 mg/kg and volume of 4 μl/g, via a cannula connected to a PE50 tube filled with saline, followed by a 6-minute equilibration period to achieve a stable hemodynamic and chronotropic steady state before data acquisition. Raw data for both conditions were reconstructed into 30 cine frames per cardiac cycle to ensure adequate temporal resolution during dobutamine-induced tachycardia.

### 2.10 Image data analysis

Quantitative image analysis was performed using Segment software (version 3.2 R8351, Medviso, Lund, Sweden). LV endocardial borders were manually traced for all slices and frames throughout the cardiac cycle. Segmentation yielded a volume–time curve for each subject; end-diastolic volume (EDV) and end-systolic volume (ESV) were defined as the global maximum and minimum of the curve, respectively. Total LV volumes were calculated using Simpson’s rule (method of disks). Functional parameters were derived as follows: stroke volume SV was computed as the difference between EDV and ESV; ejection fraction EF was expressed as the percentage ratio of SV to EDV; cardiac output CO was obtained by multiplying SV by the mean HR recorded during scanning. The peak ejection rate (PER, µl/s) and peak filling rate (PFR, µl/s) were peak absolute rates of volume change (|dV/dt|max) extracted from the first derivative of the volume–time curve, separately for the systolic and diastolic phases. To account for differences in body size, volumetric and flow parameters were normalized to body surface area BSA (cm^2^) which was derived from body weight (BW, in grams) according to the Meeh equation: BSA = 9.822 × BW^(2/3). Normalization yielded indexed parameters, including end-diastolic volume index (EDVI, µl/cm^2^), end-systolic volume index (ESVI, µl/cm^2^), stroke volume index (SVI, µl/cm^2^), and cardiac index (CI, µl/cm^2^). These measurements provided a comprehensive characterization of both inotropic and lusitropic responses to pharmacological stress.

### 2.11 Statistical data analysis

All statistical analyses were performed using GraphPad Prism 10.4.2 for Windows (GraphPad Software). For experiments assessing genotype and sex, data were analyzed using two-way ANOVA with genotype and sex as between-subject factors. Dobutamine experiments were analyzed using three-way repeated-measures ANOVA, with dobutamine condition (Pre vs. Post) as the within-subject factor and genotype and sex as between-subject factors. Where appropriate, multiple comparisons were performed using Šídák’s test. Statistical significance was set at α = 0.05. Data are presented as mean ± s.e.m.. Potential outliers were assessed using the ROUT method with Q = 1% for Body weight [g], distance moved [m] and total time in light box [s] in Figure 2, heart rate [bpm] in Figure 5, heart rate [bpm] in Pre and Post dobutamine in Figure 6, and no observations were excluded.

**Figure 2.**
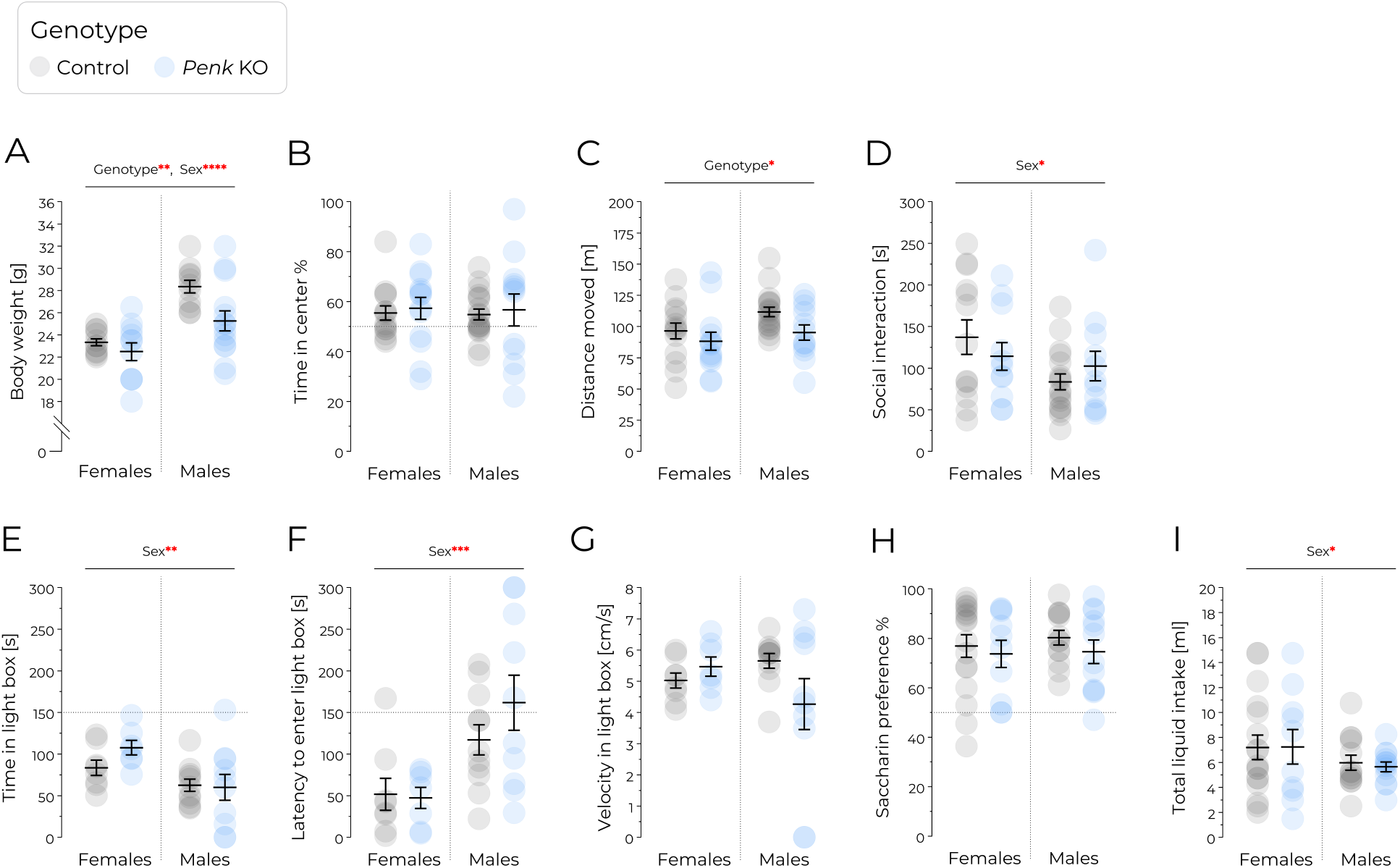
*Penk* KO mice display normal exploratory behavior, anxiety-like responses, sociability, and no anhedonia-like behavior. (A) Body weight [g]. **(B–D)** Exploratory and social behavior in the open field. (B) Time spent in the center of the open field %. (C) Total distance moved [m]. (D) Time spent in social interaction [s]. **(E–G)** Anxiety-like behavior in the light-dark box. (E) Total time spent in the light box [s]. (F) Latency to enter the light box [s]. (G) Velocity in light box [cm/s]. **(H–I)** Saccharin preference task. (H) Preference of the saccharin solution vs. water %. (I) Total liquid intake [ml]. Control and *Penk* KO animals are shown in gray and blue, respectively. Circles represent individual animals. Sample sizes were: (A) female control n = 11, female *Penk* KO n = 11, male control n = 11, male *Penk* KO n = 12; (B– C) female control n = 14, female *Penk* KO n = 13, male control n = 19, male *Penk* KO n = 12; (D) female control n = 13, female *Penk* KO n = 11, male control n = 19, male *Penk* KO n = 11; (E–G) female control n = 8, female *Penk* KO n = 7, male control n = 11, male *Penk* KO n = 10; (H–I) female control n = 17, female *Penk* KO n = 10, male control n = 13, male *Penk* KO n = 12. Data are mean ± SEM. Statistical analysis was performed using two-way ANOVA. *p<0.05, **p<0.01, ***p<0.001, ****p<0.0001.Control and *Penk* KO animals are shown in gray and blue, respectively. Circles represent individual animals. Whiskers indicate mean and s.e.m.. Statistical analysis was performed using two-way ANOVA followed by Tukey’s HSD. Significance is indicated as *p<0.05, **p<0.01, ***p<0.001, ****p<0.0001.

## 3 Results

### 3.1 A new Penk KO mouse strain and its behavioral phenotype

The new mouse strain was generated by introducing loxP sites immediately upstream of the Penk protein-coding sequence and immediately downstream of the stop codon (**Figure 1A**). Mice were crossed with a strain expressing Cre recombinase in the germ line (Syn1-cre) to remove the sequence flanked by loxP sites. PCR genotyping confirmed deletion of the targeted sequence, while qPCR showed no detectable Penk transcript in the hearts of Penk KO mice (Zenodo: 21919706). Likewise, *in situ* hybridization using RNAscope probes found no *Penk* mRNA in the brain’s striatum, where indirect pathway medium spiny neurons normally express high levels of *Penk* (**Figure 1B**). In contrast, the prodynorphin transcript remained abundantly expressed in direct pathway medium spiny neurons. Together, these observations confirm complete inactivation of the *Penk* gene in mutant mice.

The loss of *Penk* did not result in overt abnormalities. Adult body weight was significantly lower in Penk KO mice than in controls, irrespective of sex (**Figure 2A**, genotype F_1,41_ = 6.44, p = .015, genotype × sex F_1,41_ = 2.01, p = .164). As expected, females weighed less than males (**Figure 2A**, sex F_1,4_ = 28.10, p < .0001). Unlike previously reported *Penk* knockout models [25,26], *Penk KO* mice in our study showed normal exploratory behavior in a novel, non-anxiogenic setting. They spent a similar proportion of time in the center of the unlit open field as wild-type controls (**Figure 2B**, genotype F_1,53_ = .23, p = .637, sex F_1,53_ = .03, p = .875, genotype × sex F_1,53_ < .001, p = .995). However, *Penk KO* animals displayed significantly reduced locomotor activity compared to wild-type mice (**Figure 2C**, genotype F_1,53_ = 4.607, p = .036, sex F_1,53_ = 3.73, p = .058, genotype × sex F_1,53_ = .51, p = .478).

Social interaction with novel, age- and sex-matched conspecific was unaffected by genotype or the genotype × sex interaction (**Figure 2D**, genotype F_1,48_ = .01, p = .905, genotype × sex F_1,48_ = 1.71, p = .197), consistent with findings from an earlier *Penk KO* model on a C57BL/6J background [27]. Females engaged in significantly more social interaction than males (**Figure 2D**, sex F_1,48_ = 4.09, p = .049).

In the light–dark box test, *Penk* deletion did not influence anxiety-like behavior. No genotype-dependent differences were observed in time spent in the light box (**Figure 2E**, genotype F_1,32_ = 0.89, p= .352, sex F_1,32_ = 9.28, p = .005, genotype × sex F_1,32_ = 1.40, p = .245) or latency to enter the light box (**Figure 2F**, genotype F_1,32_ = .70, p = .408, sex F_1,32_ = 13.99, p = .001, genotype × sex F_1,32_ = 1.04, p = .317). Across these measures, males showed greater anxiety-like behavior than females, spending less time in the light compartment and showing longer entry latencies. Velocity in the light compartment was unaffected by genotype, sex or their interaction (**Figure 2G**, genotype F_1,32_ = .84, p = .367, sex F_1,32_ = .31, p = .581, genotype × sex F_1,32_ = 3.19, p = .084). These findings contrast with previous reported *Penk KO* strains, which showed increased anxiety-like behavior [25,26].

Anhedonia, a core symptom of depression and an independent predictor of mortality, is commonly observed in patients with heart failure, hypotension, and peripheral arterial disease. To evaluate this phenotype, we employed the saccharin preference test. *Penk KO* mice showed no differences in saccharin preference (**Figure 2H**, genotype F_1,48_ = .93, p = .340, sex F_1,48_ = .22, p = .645, genotype × sex F_1,48_ = .08, p = .778) or total liquid consumed (**Figure 2I**, genotype F_1,48_ = .02, p = .875, sex F_1,48_ = 2.36, p = .131, genotype × sex F_1,48_ = .04, p = .838) compared to controls.

Taken together, these results show that *Penk KO* mice in this study exhibit no behavioral deficits, no evidence of increased anxiety-like behaviors, and no signs of anhedonia, which contrasts with the phenotype reported in the previously generated strains [25,26].

### 3.2 Heart function in Penk KO mice

We hypothesized that inactivation of *Penk* and the loss of enkephalins would cause heart hypertrophy and may lead to other disruptions in normal heart function. However, we found no difference in the raw heart weight of *Penk* KO mice compared to controls (**Figure 3A**, genotype F_1,32_ = .19, p = .669, sex F_1,32_ = 9.69, p = .004, genotype × sex F_1,32_ = 1.54, p = .224). Likewise, there was no effect of the mutation on the weight normalized to tibia length (**Figure 3B**, genotype F_1,32_ = .02, p = .088, sex F_1,32_ = 8.99, p = .005, genotype × sex F_1,32_ = 1.80, p = .189). As expected, males had heavier hearts than females, and this effect was consistent in the raw and tibia-normalized heart weight.

**Figure 3.**
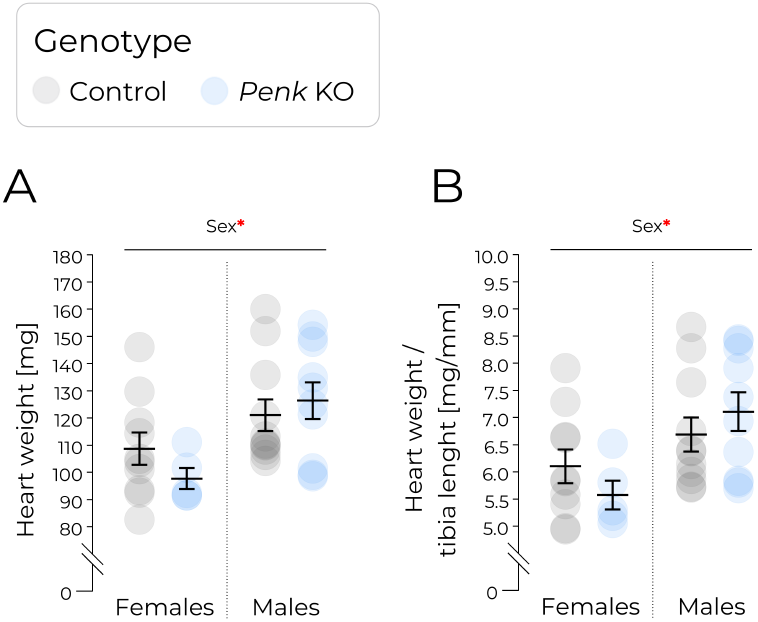
Heart weight is preserved in *Penk* KO mice. (A) Heart weight [mg]. (B) Heart weight normalized to tibia length [mg/mm]. Control and *Penk* KO animals are shown in gray and blue, respectively. Circles represent individual animals. Sample sizes were: female control n = 13, female *Penk* KO n = 11, male control n = 12, male *Penk* KO n = 14. Data are mean ± SEM. Statistical analysis was performed using two-way ANOVA. *p<0.05.

To further investigate the effect of the mutation on the heart, we have measured left ventricular function using magnetic resonance imaging in isoflurane-anesthetized mice. **Figure 4** shows representative MRI images of the ventricles from male and female control and Penk KO mice, obtained as described in the Methods section.. The analysis of MRI data showed no effect of the mutation or sex on heart rate, which was around 545 bpm in anesthetized animals (**Figure 5A**, genotype F_1,41_ = .19, p = .666, sex F_1,41_ = .34, p = .564, genotype × sex F_1,41_ = .63, p = .433). However, both the mutation and sex had a significant effect on peak fill rate (**Figure 5B**, genotype F_1,41_ = 4.13, p = .049, sex F_1,41_ = 4.42, p = .042, genotype × sex F_1,41_ = .33, p = .567), and peak ejection rate (**Figure 5C**, genotype F_1,41_ = 4.15, p = .048, sex F_1,41_ = 4.89, p = .033, genotype × sex F_1,41_ = 1.26, p = .27). Both effects reflected lower peak filling and ejection rates in *Penk* KO mice compared with controls, and in females compared with males. Nevertheless, there was no effect of the mutation on the end diastolic volume (**Figure 5D**, genotype F_1,39_ = 4.05, p = .051, sex F_1,39_ = 4.31, p = .045, genotype × sex F_1,39_ = .31, p = .576) or the end systolic volume (**Figure 5E**, genotype F_1,41_ = 1.59, p = .214, sex F_1,41_ = 5.84, p = .020, genotype × sex F_1,41_ = .26, p = .613), the two key parameters of the ventricle function. As expected, there was a significant effect of sex on both parameters, the rates were higher in females than males. Furthermore, the mutation did not affect the stroke volume index (**Figure 5F**, genotype F_1,40_ = .38, p = .541, sex F_1,40_ = .19, p = .662, genotype × sex F_1,40_ = .23, p = .628). Accordingly, the ejection fraction was also not affected by *Penk* inactivation and was significantly higher in male compared to female mice (**Figure 5G**, genotype F_1,41_ = 1.39, p = .244, sex F_1,41_ = 7.33, p = .01, genotype × sex F_1,41_ =.24, p = .624). Finally, the cardiac index did not differ between genotypes (**Figure 5H**, genotype F_1,40_ = .01, p = .909, sex F_1,40_ = 1.39, p = .244, genotype × sex F_1,40_ = .07, p = .787). Taken together, inactivation of *Penk* moderately decreased the peak ejection and filling rates, whereas other parameters of left ventricular function were not affected by the mutation.

**Figure 4.**
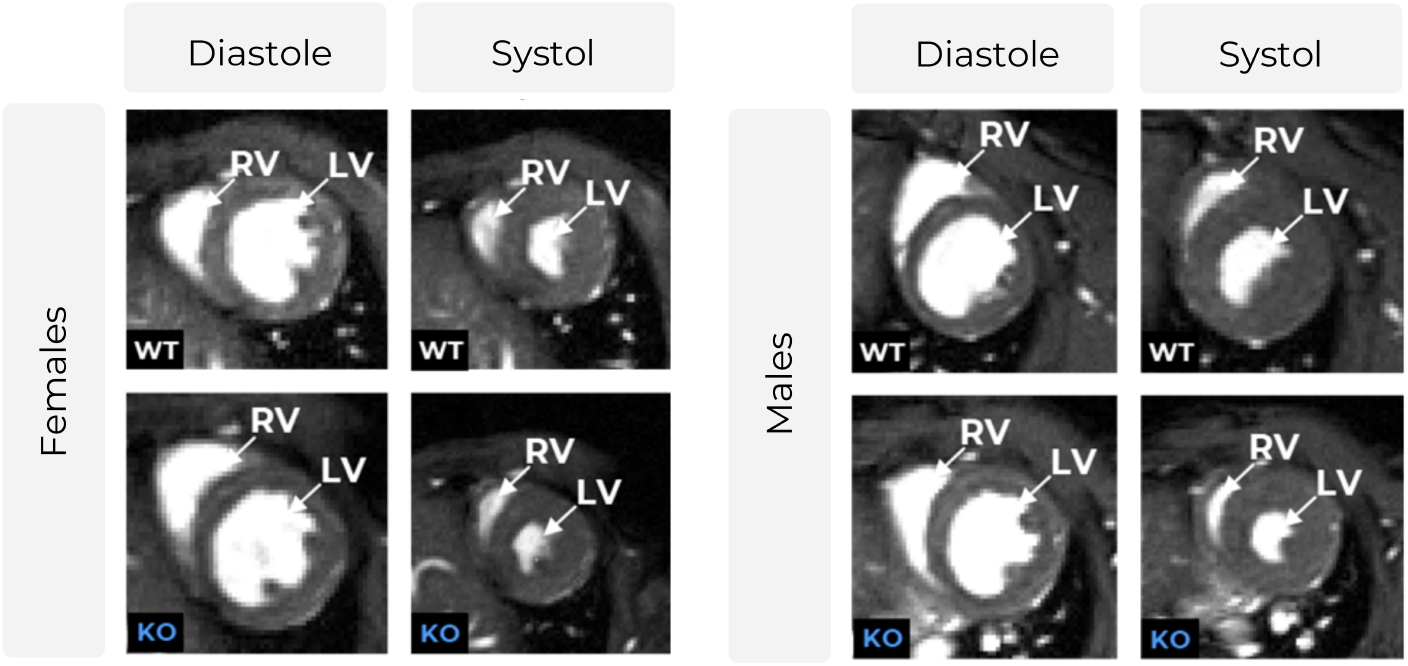
Representative cardiac MR images from control and *Penk* KO mice. Images represent a short axis myocardium slice at end-diastole and end-systole. WT – control mice, KO – *Penk* KO mice, RV – right ventricle, LV – left ventricle. **O**ne representative animal from each indicated group is displayed.

**Figure 5.**
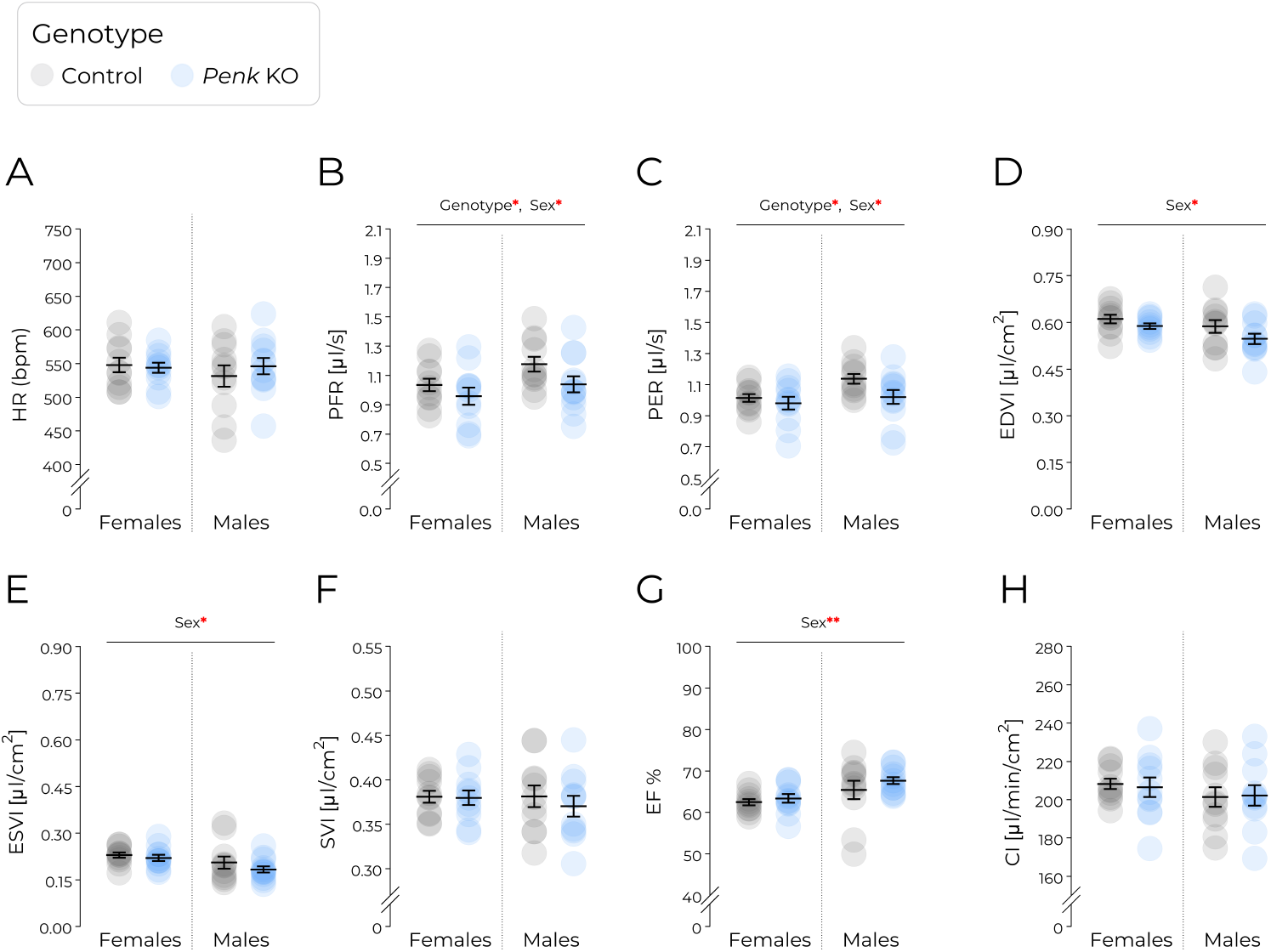
Basal left ventricular function in control and *Penk* KO mice. (A) Heart rate (HR, bpm). (B) Peak filling rate (PFR, [μl/s]). (C) Peak ejection rate (PER, [μl/s]). Body surface area (BSA) = 9.822 x body weight^2/3 [cm2]. (D) End-diastolic volume index divided by BSA (EDVI, [μl/cm^2^]). (E) End-systolic volume index divided by BSA (ESVI, [μl/cm^2^]). (F) Stroke volume divided by BSA (SVI, [μl/cm^2^]). (G) Ejection fraction (EF, %). (H) Cardiac index (CI, [μl/min/cm^2^]). Control and *Penk* KO animals are shown in gray and blue, respectively. Circles represent individual animals. Sample sizes were: female control n = 11, female *Penk* KO n = 11, male control n = 11, male *Penk* KO n = 12. Data are mean ± SEM. Statistical analysis was performed using two-way ANOVA. *p<0.05, **p<0.01.

Next, we assessed left ventricular function following stimulation with the β_1_-adrenergic agonist dobutamine. As expected, dobutamine treatment significantly increased heart rate with no significant effects of sex, genotype, or their interactions (**Figure 6A**, dobutamine F_1,32_ = 34.07, p < .0001, genotype F_1,32_ = .81, p = .374, sex F_1,32_ = .17, p = .683, dobutamine x sex F_1,32_ = .01, p = .939, dobutamine x genotype, F_1, 32_ = .60, p = .444, sex x genotype, F_1, 32_ = .24, p = .629, dobutamine × sex × genotype F_1, 32_ = .71, p = .405). It also reduced the interval between heartbeats. There was a significant interaction between dobutamine and sex on the peak filling rate (**Figure 6B**, dobutamine F_1,40_ = 9.68, p = .003, sex F_1,40_ = 4.21, p = .047, genotype F_1,40_ = .01, p = .913, dobutamine × sex F_1,40_ = 4.44, p = .041, dobutamine × genotype, F_1,40_ = 1.88, p = .178, sex × genotype, F_1,40_ = .01, p = .913, dobutamine × sex × genotype F_1,40_ =

**Figure 6.**
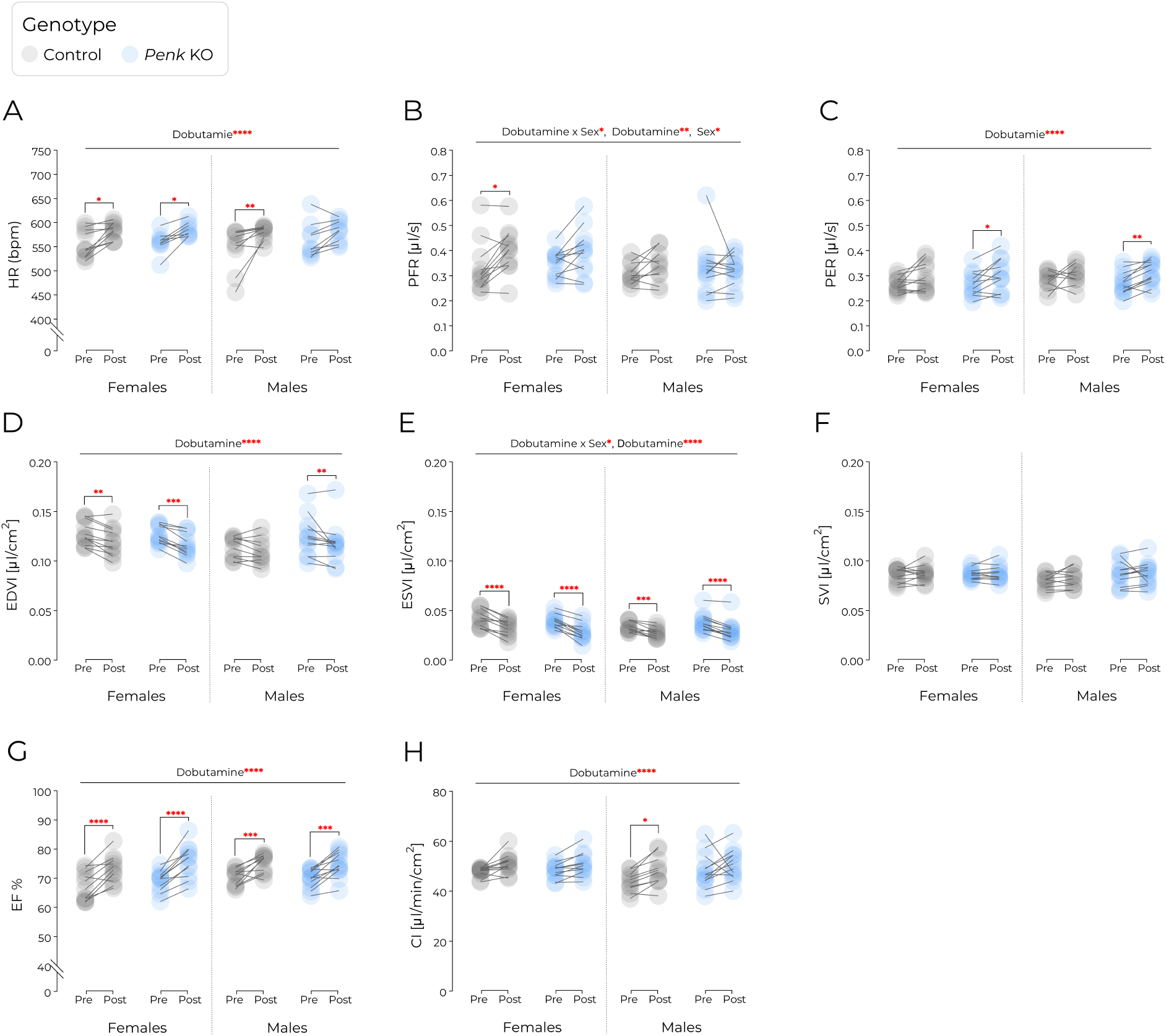
Left ventricular response to dobutamine is preserved in Penk KO mice. (A) Heart rate (HR, bpm). (B) Peak filling rate (PFR, [μl/s]). (C) Peak ejection rate (PER, [μl/s]). Body surface area (BSA) = 9.822 x body weight^2/3 [cm2]. (D) End-diastolic volume index divided by BSA (EDVI, [μl/cm^2^]). (E) End-systolic volume index divided by BSA (ESVI, [μl/cm^2^]). (F) Stroke volume divided by BSA (SVI, [μl/cm^2^]). (G) Ejection fraction (EF, %). (H) Cardiac index (CI, [μl/min/cm^2^]). Data represents parameters before (Pre) and after (Post) dobutamine administration. Control and *Penk* KO animals are shown in gray and blue, respectively. Each pair of connected observations represents repeated measurements from the same animal. Sample sizes were: female control n = 11, female *Penk* KO n = 11, male control n = 11, male *Penk* KO n = 12. Data are mean ± SEM. Statistical analysis: three-way ANOVA with dobutamine condition (Pre/Post) as the within-subject factor and sex and genotype as between-subject factors followed by Šídák’s multiple-comparisons test. Significance is indicated as *p<0.05, **p<0.01, ***p<0.001, ****p<0.0001.

.16, p = .688). Females showed a greater increase in peak filling rate than males, while genotype had no significant effect. Dobutamine also robustly increased the peak ejection rate without significant effects of sex, genotype, or interactions between factors (**Figure 6C**, dobutamine F_1,40_ = 28.83, p < .0001, sex F_1,40_ = .585, p = .449, genotype F_1,40_ = .012, p = .918, dobutamine × sex F_1,40_ = .066, p = .799, dobutamine × genotype, F_1,40_ = 1.179, p = .284, sex x genotype, F_1,40_ = .12, p = .730, dobutamine × sex × genotype F_1,40_ = .33, p = .567). Treatment decreased both end-diastolic volume (**Figure 6D**, dobutamine F_1,40_ = 36.36, p < .0001, sex F_1,40_ = 2.06, p = .156, genotype F_1,40_ = .59, p = .445, dobutamine x sex F_1,40_ = 2.44, p = .126, dobutamine x genotype, F_1,40_ = 2.048, p = .160, sex × genotype, F_1,40_ = 1.93, p = .171, dobutamine × sex × genotype F_1,40_ = .30, p = .586) and systolic volume (**Figure 6E**, dobutamine F_1,40_ = 151.2, p < .0001, sex F_1,40_ = 2.21, p = .145, genotype F_1,40_ = .01, p = .941, dobutamine x genotype, F_1,40_ = .90, p = .350, sex × genotype, F_1,40_ = 2.11, p = .154, dobutamine × sex × genotype F_1,40_ = .03, p = .866). There was a significant dobutamine × sex interaction for systolic volume (dobutamine x sex F_1,40_ = 6.15, p = .017), indicating a larger decrease in females. Genotype did not influence either parameter. The stroke volume index was not significantly affected by dobutamine treatment, sex, or genotype (**Figure 6F**, dobutamine F_1,40_ = 1.23, p = .274, sex F_1,40_ = 1.07, p = .307, genotype F_1,40_ = 1.83, p = .184, dobutamine x sex F_1,40_ = .01, p = .889, dobutamine x genotype, F_1,40_ = .86, p = .358, sex x genotype, F_1,40_ = .99, p = .326, dobutamine x sex x genotype F_1,40_ = .23, p = .636). In contrast, dobutamine significantly increased the ejection fraction (**Figure 6G**, dobutamine F_1,40_ = 102.0, p < .0001, sex F_1,40_ = 1.37, p = .248, genotype F_1,40_ = .70, p = .406, dobutamine x sex F_1,40_ = 2.25, p = .141, dobutamine x genotype, F_1,40_ = .11, p = .740, sex x genotype, F_1,40_ = 1.25, p = .271, dobutamine x sex x genotype F_1,40_ = .10, p = .749) and cardiac index (**Figure 6H**, dobutamine F_1,40_ = 19.87, p < .0001, sex F_1,40_ = 1.75, p = .194, genotype F_1,40_ = 1.62, p = .210, dobutamine x sex F_1,40_ = .166, p = .686, dobutamine x genotype, F_1,40_ = 1.29, p = .262, sex x genotype, F_1,40_ = 1.31, p = .258, dobutamine x sex x genotype F_1,40_ = .25, p = .621), again with no effects of sex, genotype, or interactions. Taken together, these data indicate that *Penk* inactivation does not alter cardiac responses to dobutamine stimulation.

## 4 Discussion

We find that the loss of *Penk*-derived enkephalins causes no overt behavioral phenotype, no appreciable heart hypertrophy in adult animals, and only a modest alteration in heart function. The left ventricle peak ejection and fill rate were significantly lower in *Penk* KO mice. However, there were no mutation effects on the end diastolic and systolic volumes, and Penk KO mice exhibited normal increases in left ventricular activity after dobutamine treatment.

This study was performed on a novel Penk KO strain, which, unlike the two previously generated mutations, has a complete deletion of the entire coding sequence [25,26,28]. Surprisingly, unlike previously reported Penk KO strains, we observed no significant changes in anxiety-like behaviors. An increase in anxiety would be intuitively consistent with the proposed roles of brain enkephalins [29], and the observed anxiolytic effects of δ receptor agonists e.g. [30,31]. We have no definitive explanation for this discrepancy. It is possible that the experiments presented here were conducted under minimal stress conditions, with the animals always group-housed in an enriched environment, capturing an unchanged baseline of anxiety-like behaviors.

We find that the hearts of adult male and female Penk KO mice are not significantly different in size from those of controls. Previously, it was reported that treatment with naltrexone during pregnancy in rats caused an increase in heart weight of the offspring that persisted in females at least till 75 days of age [21]. This suggests that in mice, the absence of ζ-opioid receptor activation can be compensated for by other mechanisms to achieve normal adult heart dimensions, or that the receptor regulates growth rate rather than the final size. Furthermore, there was no effect of mutation on basal or sympathomimetic-stimulated heart contractility or heart rate, despite the reported continuous release of enkephalins in the heart [7]. Most importantly, the left ventricular function was only moderately affected by *Penk* KO, and key parameters remained normal. Lower peak ejection and fill rates might theoretically limit performance under high heart activity, but this remains speculative. Still, we argue that our data show that *Penk* is not essential for heart development or normal physiology. Some reports indicate that *Penk* expression increases with age [14,32] or in response to heart disease [33], suggesting a potential role in older animals, whereas our study focused on adults. Further research could explore long-term effects of stress or ageing on cardiac function in the absence of enkephalin signaling.

The study was conducted in both male and female mice, allowing observation of sex-dependent differences. We found that females exhibited longer social interaction times than males, regardless of genotype. Shorter interaction times are typically associated with increased social anxiety [34]. Accordingly, females also showed reduced anxiety-like behavior, spending more time in the light box, with shorter latency to enter the compartment and greater distance moved. This suggests that lower baseline anxiety may underlie their increased social interaction, consistent with previous literature [35]. Although human females often outperform males in social domains [36,37], evidence for similar sex differences in rodents is inconsistent [38,39]. Prior studies using social interaction in open field often reported greater social engagement in male mice [40,41], though females sometimes showed increased social contact-seeking depending on context [42]. No sex differences were observed in open field center time or distance moved, consistent with this measure reflecting exploratory behavior in a low-anxiety environment, highlighting the importance of context when interpreting sex differences in social and anxiety-related behaviors.

In conclusion, we find that a complete inactivation of the *Penk* gene produces only mild alterations in heart function and does not cause overt behavioral deficits in adult mice.

## CRediT authorship contribution statement

Klaudia Misiołek: investigation, data curation, methodology, visualization, writing – original draft, writing – review and editing; Łukasz Szumiec: investigation, writing – review and editing, Krzysztof Jasiński: investigation, formal analysis, methodology, writing – review and editing; Katarzyna Kalita: investigation, methodology, writing – review and editing; Maria Kaczmarzyk-Jarosz: investigation, writing – review and editing; Magdalena Chrószcz: investigation, writing – review and editing; Władysław P. Węglarz, conceptualization, formal analysis, supervision, writing – original draft, writing – review and editing; Barbara Ziółkowska: conceptualization, supervision, writing – original draft, writing – review and editing; Jan Rodriguez Parkitna: conceptualization, funding acquisition, supervision, writing – original draft, writing – review and editing.

## Data availability

All results used to prepare the figures are available on Zenodo at **10.5281/zenodo.21919706**. Raw data from behavioral video recordings and raw MRI data are available from corresponding author upon reasonable request.

