## Supplementary data for "The effect of preproenkephalin gene deletion on heart function in mice"

| **Supplementary Table S1.** Animals used in Figures 1 and 3-6*. | | | | | | | | | | |
| --- | --- | --- | --- | --- | --- | --- | --- | --- | --- | --- |
| **Experiment** | **Sex** | | **Genotype** | **Animals tested (n)** | **Subject's weight [g]** | |  | **Subject's age (weeks)** | | **Outliers (n)** |
|  |  |  |  |  | **Mean ± SEM** | **Range** |  | **Mean ± SEM** | **Range** |  |
| **qPCR** (Fig. 1) and **Heart weight** (Fig. 3 | | Females | Control | 13 | 20.7 ± 0.3 | 4 |  | 22.0 ± 0.7 | 6 | - |
|  |  |  | *Penk KO* | 11 | 20.1 ± 1.1 | 3 |  | 21.4 ± 0.6 | 8 | - |
|  |  | Males | Control | 12 | 24.4 ± 0.8 | 9 |  | 20.5 ± 0.8 | 9 | - |
|  |  |  | *Penk KO* | 14 | 25.1 ± 0.5 | 7 |  | 21.9 ± 0.6 | 9 | - |
| **Body weight** (Fig. 2A) and **MRI** (Fig. 4-6) | | Females | Control | 11 | 23.3 ± 0.3 | 3 |  | 22.0 ± 1.4 | 13 | - |
|  |  |  | *Penk KO* | 11 | 22.5 ± 0.8 | 8 |  | 21.2 ± 1.4 | 13 | - |
|  |  | Males | Control | 11 | 28.4 ± 0.6 | 6 |  | 24.1 ± 0.9 | 8 | - |
|  |  |  | *Penk KO* | 12 | 25.4 ± 0.6 | 11 |  | 20.4 ± 2.0 | 20 | - |
| *A subset of mice (n = 4) from heart weight experiment was used for qPCR analysis. Representative images from animals included in MRI experiment are shown in Figure 4. No data points were identified as outliers by ROUT test applied to: Body weight [g], HeartRate [bpm] in Figure 5, HeartRate [bpm] in Pre and Post dobutamine in Figure 6. | | | | | | | | | | |
| **Supplementary Table S2.** Animals used in Figure 2*. | | | | | | | | | | |
| **Experiment** | **Sex** | | **Genotype** | **Animals tested (n)** | **Subject's weight [g]** | |  | **Subject's age (weeks)** | | **Outliers (n)** |
|  |  |  |  |  | **Mean ± SEM** | **Range** |  | **Mean ± SEM** | **Range** |  |
| Open field | | Females | Control | 14 | 20.7 ± 0.3 | 4 |  | 11.1 ± 0.5 | 6 | - |
|  |  |  | *Penk KO* | 13 | 19.5 ± 0.6 | 6 |  | 11.8 ± 0.6 | 7 | - |
|  |  | Males | Control | 19 | 24.4 ± 0.5 | 9 |  | 12.6 ± 0.4 | 5 | - |
|  |  |  | *Penk KO* | 12 | 24.3 ± 0.6 | 7 |  | 11.8 ± 0.4 | 5 | - |
| Social interaction | | Females | Control | 13 | 20.3 ± 0.3 | 4 |  | 12.1 ± 0.5 | 5 | - |
|  |  |  | *Penk KO* | 11 | 19.2 ± 0.6 | 6 |  | 12.7 ± 0.4 | 3 | - |
|  |  | Males | Control | 19 | 24.4 ± 0.5 | 9 |  | 12.6 ± 0.4 | 5 | - |
|  |  |  | *Penk KO* | 11 | 24.3 ± 0.7 | 7 |  | 11.9 ± 0.4 | 5 | - |
| Light-dark box | | Females | Control | 8 | 17.8 ± 0.3 | 3 |  | 15.9 ± 1.1 | 8 | - |
|  |  |  | *Penk KO* | 7 | 17.3 ± 0.4 | 3 |  | 13.1 ± 0.3 | 3 | - |
|  |  | Males | Control | 11 | 26.2 ± 0.7 | 7 |  | 18.4 ± 1.0 | 8 | - |
|  |  |  | *Penk KO* | 10 | 22.8 ± 0.4 | 4 |  | 14.2 ± 0.8 | 6 | - |
| Saccharin preference | | Females | Control | 17 | NA | NA |  | 14.4 ± 0.9 | 13 | - |
|  |  |  | *Penk KO* | 10 | NA | NA |  | 13.8 ± 0.2 | 2 | - |
|  |  | Males | Control | 13 | NA | NA |  | 18.6 ± 1.2 | 9 | - |
|  |  |  | *Penk KO* | 12 | NA | NA |  | 14.2 ± 0.8 | 9 | - |
| *Except for Figure 2A, which presents body weight data from animals included in the MRI experiments shown in Figures 5 and 6. No data points were identified as outliers by the ROUT test applied to: OF = distance moved, LDB = total time in light box. | | | | | | | | | | |
